# The fine-tuning of synapse development by oxidative stress and autophagy requires presynaptic ATM kinase

**DOI:** 10.1101/2024.04.25.591136

**Authors:** Matthew J. Taylor, Syed Azan Ahmed, Ellena G. Badenoch, David Bennett, Richard I. Tuxworth

**Affiliations:** Institute of Cancer and Genomic Sciences, University of Birmingham, Birmingham B15 2TT, UK

**Keywords:** ATM, ROS, autophagy, ataxia-telangiectasia, *Drosophila*, neurodegeneration

## Abstract

Two processes held in delicate balance during the fine tuning of synapse development are oxidative stress and autophagy: each can promote synapse expansion yet in excess are toxic. How this balance is maintained is not fully understood. While ataxia-telangiectasia mutated (ATM) is recognized as a key regulator of the DNA damage response, there is increasing evidence of a neuronal-specific role for this ubiquitous kinase and deficiency causes early-onset neurodegeneration. We report a requirement for presynaptic *Drosophila* ATM (dATM) in neurodevelopment that is independent of its functions in the DNA damage response. Reduction of presynaptic *dATM* expression causes hypersensitivity to raised oxidative stress and a failure to induce autophagy which leaves mitochondria in excess in neurons. We demonstrate that presynaptic dATM coordinates autophagy through the conserved ATM-AMPK axis. Similarly to mammalian ATM, neuronal dATM is predominantly cytosolic and forms synaptic foci. dATM also colocalizes with autophagosomes. We propose a model wherein dATM responds to increased reactive oxygen species resulting from heightened neuronal activity by activating autophagy to induce synaptic growth, while protecting the neuron from excitotoxicity and oxidative stress.

## Introduction

As synapses mature and change over time and with new experiences, the neuron must integrate different signals which, in other contexts, can be toxic. For example, oxidative species such as hydrogen peroxide can be used by neurons at physiological levels as an instructive signal reporting increased presynaptic activity levels to regulate synapse homeostasis, specifically increasing presynaptic arborisation and modulating synaptic output (1,2). However, at the upper end of its physiological concentration range, hydrogen peroxide is toxic and causes neurodegeneration (3). A similar story is true for macroautophagy (referred to here as autophagy), the subcellular recycling programme which digests damaged cellular components and organelles. Autophagy has been identified as necessary for normal expansion of synapses during development (4) but, like oxidative stress, an excess of autophagy can be deleterious (5). We do not fully understand the mechanisms through which neurons balance the competing need for oxidative stress and autophagy with the threat of toxicity when in excess. The consequences when this balancing act goes awry can be observed in numerous neurodegenerative disorders, including Alzheimer’s, Parkinson’s and Huntington’s diseases (6–10).

Ataxia-telangiectasia (A-T), which is caused by mutations in ATM kinase, is an early-onset neurodegenerative disorder in which the cerebellum is particularly vulnerable – although it is still an open question as to why (11–13). ATM kinase is a key regulator of the DNA damage response (DDR) but has other, non-nuclear functions that depend upon both its subcellular localisation and mechanism of activation, including stimulation of autophagy and mitophagy (macroautophagy of mitochondria) in the cytosol following activation of dimeric ATM by reactive oxygen species, which activate the kinase activity (14–16). There is increasing evidence of a neuronal-specific role for the ATM protein unrelated to its role in the DDR. From early studies of this kinase, it was noticed that neurons have a particularly large pool of cytosolic ATM compared to other cell types (17–19) and a subset of ATM co-localises with presynaptic vesicles in synapses and is required for long-term potentiation (20).

In A-T, various lines of evidence ranging from mouse models to human cell lines indicate autophagic flux is altered as a consequence of ATM deficiency (21–23). In addition, ATM deficient cells are vulnerable to increased oxidative stress (24). Together, these point to a potential role for ATM regulating how synapses respond to these instructive signals and that the neurodegeneration in A-T may be due to a failure to balance the risk of toxicity they bring.

Rodent models of A-T recapitulate the immunological, fertility and radiosensitivity aspects of the disease, with some evidence of cerebellar dysfunction, such as poorer performance on the rota-rod test or wider foot spacing in gait analysis tests (25–27). However, mouse models of A-T do not show evidence of cerebellar neurodegeneration – a hallmark of the human disease – or other neurodevelopmental defects (26–28).

We sought to define a role for ATM in synapse development and homeostasis using the well-characterised model of the *Drosophila* larval neuromuscular junction (NMJ). We find that presynaptic ATM is required for normal synapse development and function. Neurons depleted of ATM fail to induce autophagy via activation of AMP kinase, resulting in a failure to develop to the correct size. ATM-depleted neurons are on the cusp of neurodegeneration and hypersensitive to a rise in oxidative stress. Our data suggest that presynaptic ATM is necessary for neurons to balance the positive, pleiotropic effects of autophagy and reactive oxidative species with their inherent toxicity when in excess.

## Methods

### *Drosophila* stocks and maintenance

Experimental crosses were reared on standard yeast-glucose-agar food using a 12 h:12 h light:dark cycle at 25°C throughout egg-laying and larval development unless otherwise stated. The dual-colour autophagy reporter *UAS-GFP-Cherry::Atg8a* was a kind gift of I. Nezis, University of Warwick, UK. *UAS-dATM[msGFP2]* was generated in this study through synthesis of the full-length *dATM* cDNA containing codon-optimised *msGFP2* (GenScript, USA), sub cloning into *pUAST-attB* and injection into the *attP-3B* landing site on chromosome 2 (BestGene, USA). The following were sourced from the Bloomington *Drosophila* Stock Centre (BDSC): *w^1118^* (BDSC #5905); *dATM^-3^* (BDSC #8625); *dATM^-6^* (BDSC #8626); *dATM^-8^* (BDSC #8624); *elav-GAL4* (BDSC #25750); *mef2-GAL4* (BDSC #27390); *repo-GAL4* (BDSC #7415); *OK371-GAL4* (BDSC #26160); *dATM[TRiP.HMS02790]* (BDSC #44073); *dATM[TRiP.JF01422]* (BDSC #31635); d*MRE11[TRiP.HMC02995]* (BDSC #50628); *dATR[TRiP.HMS02331]* (BDSC #41934); *dCHK2[TRiP.HMC05499]* (BDSC #64482); *cat[TRiP.JF02173]* (BDSC #31894); *Atg18[TRiP.HMS01193]* (BDSC #34714); *AMPK[TRiP.JF01951]* (BDSC #25931); *UAS-ATG1* (BDSC #51654 and #51655); *UAS-AMPK* (BDSC #32108); *UAS-GC3Ai* (BDSC #84301 and #84313); *UAS-TrpA1* (BDSC #23263 and #26264).

### 3rd instar larval dissection

*NMJs* - *Drosophila* wandering 3rd instar-stage larval NMJ “fillet” dissections were performed as described elsewhere (29). The CNS was left intact, except for electrophysiology experiments in which it was removed to prevent spontaneous muscle contractions. The samples were fixed in 4% paraformaldehyde/PBS followed by washing in PBS prior to immunostaining.

*CNS and salivary glands* - the CNS was exposed from wandering 3^rd^ instar larvae by gentle application of tension to the mouth-hooks. The salivary glands, eye discs and any excess tissue was then separated from the CNS. The isolated CNS and salivary glands were fixed then washed in PBS prior to immunostaining or live imaging.

*For irradiation experiments*, larvae were irradiated with 8 Gy (CellRad X-ray irradiator) followed by 30 min of recovery prior to dissection.

All dissections were performed in low Ca^++^ HL3.1, an isotonic buffer that mimics the larval haemolymph (30).

### Immunohistochemistry

Dissected tissues were permeabilised for 15 min in PBT (PBS + 0.1 % v/v Triton-X100) and blocked in 1 % BSA/PBS for 1 h. The primary antibody step was performed at 4°C for 1-3 days in blocking solution, before washing in PBS. Primary antibodies were as follows: mouse anti-BRP (1:100, Developmental Studies Hybridoma Bank, University of Iowa [nc82]); chicken anti-GFP (1:400, Invitrogen, Cat# A10262); mouse anti-DLG (1:200, DSHB [4F3]); Alexa-594 goat anti-HRP (1:400, Jackson Immuno, Cat# 123-585-021). Samples were incubated in secondary antibodies in PBS 4 hr-overnight at 4°C. Secondary antibodies included: Alexa-488 donkey anti-mouse (1:1000, Jackson Immuno, Cat# 715-545-150); Alexa-488 anti-chicken (1:400, Thermo, Cat# A32931); ToPro3 (1:2000, Invitrogen, Cat# T3605). Finally, the preparations were washed in PBS and mounted on glass slides in either Fluoromount (with DAPI) or Prolong Gold (without DAPI).

### Electrophysiology

3rd instar larval electrophysiology was performed as described elsewhere (31). Larvae were dissected as for the larval NMJ preps in low Ca^++^ HL3.1 to minimise muscle contraction during dissection. The motor neuron axons were severed at the base of the CNS, and the CNS was removed to prevent spontaneous muscle contraction during recordings. The samples were washed in HL3.1 and recordings were performed in HL3.1 containing 1.5 mM Ca^++^. Recordings were taken from muscle 6/7 in segments A2-A5. Single-electrode current clamp recordings were performed using the bridge mode of an AxoClamp-2B amplifier with a HS-2A headstage (Axon Instruments), with stimulation provided by a DS2A Isolated Voltage stimulator (Digitimer Ltd).

For the recording electrode, borosilicate glass capillaries (GC150F-10, Harvard Apparatus) were pulled using a Narishige PC-100 to a final resistance of 15-25 MΩ and filled with 3 M KCl. The stimulation electrode holder was constructed following a standard protocol (32). Stimulation (suction) electrodes were made to a final resistance of 5 MΩ, the tip gently broken against a microscope lens tissue, and backfilled with HL3.1 using negative suction.

The motor neuron innervating the respective segment was identified and gentle application of negative suction brought the severed axon end into the stimulation electrode. After insertion of the recording electrode, various exclusion criteria were looked for:

- A voltage drop to a V_m_ of at least −60 mV
- A muscle R_in_ of at least 4 MΩ as measured by the voltage drop following a −1 nA current injection.
- Correctly identified segmental motor neuron – validated by manually stimulating to check that an excitatory junction potential (EJP) was evoked.
- Recruitment of both Ib and Is motor neuron inputs (see below).

EJPs were evoked initially by 200 μs stimulation at increasing voltages (ranging from 1-8 V) to find the minimum voltage required to recruit both Ib and Is motor neuron consistently, which could be seen by first a small EJP response at one threshold and then a distinct, discrete increase with addition of extra voltage. Mean EJP amplitude was calculated from 10 evoked EJPs at 0.5 Hz. Mini excitatory junction potentials (mEJPs) were observed by recording fluctuations in Vm for at least 2 minutes post-stimulation.

Data were recorded in Spike2 v9.16.

### Larval locomotion assay

Individual wandering third instar larvae were transferred into a custom-made 3D printed arena with wells containing 2% agarose coloured with a small amount of Orange-G dye. Larval locomotion while freely crawling was tracked for 5 minutes using EthovisionXT software. 8-12 larvae were recorded simultaneously. Only larvae with a mean speed and percentage time moving > 0 were used for subsequent analysis.

### Confocal microscopy

All images were taken using either a Zeiss LSM 780 or LSM 880 confocal microscope. For muscle size quantification, images were obtained using a 10X water immersion lens, using a single scan capturing at both 488 nm and 594 nm. For NMJ quantification, the NMJ on muscle 4 was imaged using a 63X water immersion lens; the images consisted of Z-stacks through the entire NMJ with 0.25 μm step size. For dATM[sfGFP] and dATM[msGFP2] localisation, NMJ images were captured using a 100X oil immersion N.A. 1.46 lens. For NMJ analysis, the resultant Zeiss raw (.czi) images were imported into FIJI for further analysis (see below).

For live imaging of the GFP-mCherry-Atg8a reporter, dissected tissues were maintained on ice in HL3.1 solution and transferred to 35 mm glass-bottom dishes. CNS and salivary gland preparations were allowed to sink to the bottom these dishes prior to live imaging, which was performed on the inverted LSM 880 microscope. CNS and salivary glands were imaged using a 25X oil immersion lens, with a single track using both the 488 nm and 594 nm laser lines, with a z-step size of 2 μm.

### Analysis of NMJ images in FIJI

Batch processing of larval NMJ images was performed using the *Drosophila* NMJ morphometrics plugin in FIJI (33). The surface area of muscle 4 was measured for each larva using the polygon selection tool and inbuilt measure function in FIJI. The mean muscle surface area (MSA) was then calculated for each genotype and the ratio of this MSA to the control MSA for each experiment generated. All datapoints were then scaled using this ratio to account for differences in muscle size between the genotypes.

### Autophagy quantification

For quantification of autophagic flux using the GFP-mCherry-Atg8a reporter (34), z-stack maximum intensity projections were analysed using a custom-made FIJI script. Essentially, the script would: iterate through each file in a directory; prompt the user to draw a freehand selection around the salivary glands; clear outside the selection; split the channels; process the GFP channel by thresholding using the auto threshold mean function and selecting the salivary gland border as the region of interest (ROI); apply this ROI to the mCherry channel; automatically adjust contrast to a fixed saturation value (to ensure consistency between samples); auto threshold (using otsu); and finally analyse particles again to select and quantify the mCherry foci. The results were exported as an Excel file and imported into Rstudio for statistical testing.

### Calculation of HRP-Dlg ratio

A custom FIJI script was written which would iterate through max intensity projections in a directory, select the HRP channel and ask the user to roughly draw a ROI around the NMJ. Auto threshold was used to detect the NMJ outline, which was then used to automatically select the entire NMJ as a ROI. This ROI was re-applied to both the HRP and DLG channels, the mean intensity of the signal measured, and the ratio of the two calculated.

### Processing of electrophysiology data

Raw electrophysiology data from Spike2 v9 were analysed using custom active cursors. For detection and quantification of EJPs, one cursor detected the points automatically marked where each stimulus was delivered, a second found the maximum value within one second of this event, while a third found the minimum. The max V_m_, min V_m_ and difference was measured.

For mEJP detection and quantification, an automatic detection pipeline was set up. The data channel was duplicated, a low pass filter applied, and a DC removal filter applied with a time constant of 50 ms. Finally, a smoothing filter with a time constant of 0.65 ms was applied to the duplicated data channel to remove high frequency noise. Active cursors were then utilised in a similar way as above, except the first cursor was set up to find peaks of at least 0.3 mV in amplitude in the DC-removed memory channel. The next cursor looked for the maximum V_m_ value within +/− 5 ms seconds of the original while the final cursor found the minimum V_m_ value within 20 ms prior to the former.

Frequency of mEJPs was calculated by taking the number of automatically detected mEJPs and dividing by the time difference between the first and last observed mEJP. Amplitudes were corrected for differences in baseline V_m_ (i.e., corrections for non-linear summation) using a derivation of Martin’s relationship (35–37):

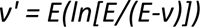

where:

- *v’* = corrected amplitude
- *E* = driving force (assumed to be equal to V_m_ given a reversal potential of 0 mV)
- *v* = recorded amplitude

Quantal content was calculated by dividing corrected EJP amplitude by corrected mEJP amplitude.

### Statistical analysis in Rstudio

All statistical analysis and graph production was performed in Rstudio v3.6.0, using the following libraries: *readxl*, *ggplot2*, *dplyr*, *ggthemes*, *ggpubr*, *ggsignif*, *ggthemr*, *tidyverse*, *ISLR*, *Rfast*, *rstatix*, *ggtext*, *RolorBrewer*, *ggsci* and *MASS*.

An R script was made for each experiment type e.g., NMJ structural analysis, electrophysiology, autophagy quantification etc. If more than one datapoint was generated from one larva (i.e., the right-side NMJ vs the left-side NMJ), then the mean of the data for that larva was used to avoid inflating the n number with non-independent datapoints.

Boxplots were generated using *ggplot2* and statistical tests performed using functions within the *rstatix* and *multcomp* packages. Data were checked for normality using the ‘shapiro.test()’ function. Student’s T-tests (with Welch’s correction) were used for pairwise comparisons of normally distributed data (Wilcoxon tests if not normally distributed), while multiple comparisons with the control genotype as the reference group were performed using Dunnett’s tests. If no group was selected as control (i.e., testing every condition against every other condition) then Tukey’s Honest Significant Differences (parametric data) or Dunn’s tests (non-parametric data) were performed. Boxplots show individual data points. Boxes represent median plus interquartile range (IQR). Whiskers represent 1.75x the IQR.

## Results

### Presynaptic dATM is required for NMJ development, expansion and function

A-T is a loss-of-function disease and is classified as a neurodevelopmental and/or early-onset neurodegenerative disorder. The early-onset neurodegeneration in A-T may well be underpinned by defective neurodevelopment yet the role ATM plays in the development of the nervous system is not well described. Our recent work indicates that ATM may have a different role in the mature nervous system of adults, where depleting it from neurons can be protective in neurological disease models (38), than it does during development. *Drosophila* is an ideal system to assess how ATM might function in neural development at the molecular level, particularly if the glutamatergic larval neuromuscular junction (NMJ) is used as the model, since it is amenable to a combination of genetic manipulation, high resolution microscopy and functional assays. Previous studies of *Drosophila* ATM (dATM) have focused predominantly on phenotypes in adult flies, such as a rough-eye phenotype or increased vacuolization of brain sections (39,40). A role for dATM in neural development remains unexplored.

We started by asking if there are neurodevelopmental deficits in whole animal *dATM* mutants. *dATM* mutants are homozygous lethal but heterozygotes are viable and fertile. The morphology of type Ib NMJ of body wall muscle 4 were quantified at the wandering 3^rd^ instar stage from confocal images after fixation and immunostaining. Significant reductions in NMJ size, active zone number, and bouton count were observed in *dATM*-/+ mutant larvae compared to controls (Fig 1A), indicating that *dATM* is haploinsufficient for NMJ development. Introducing a 20 kb BAC encompassing the *dATM* locus into the *dATM^-8/+^* background restored NMJ surface area and bouton count, confirming that the phenotype was due to haploinsufficiency of *dATM,* although active zone number was not fully restored.

**Figure 1.**
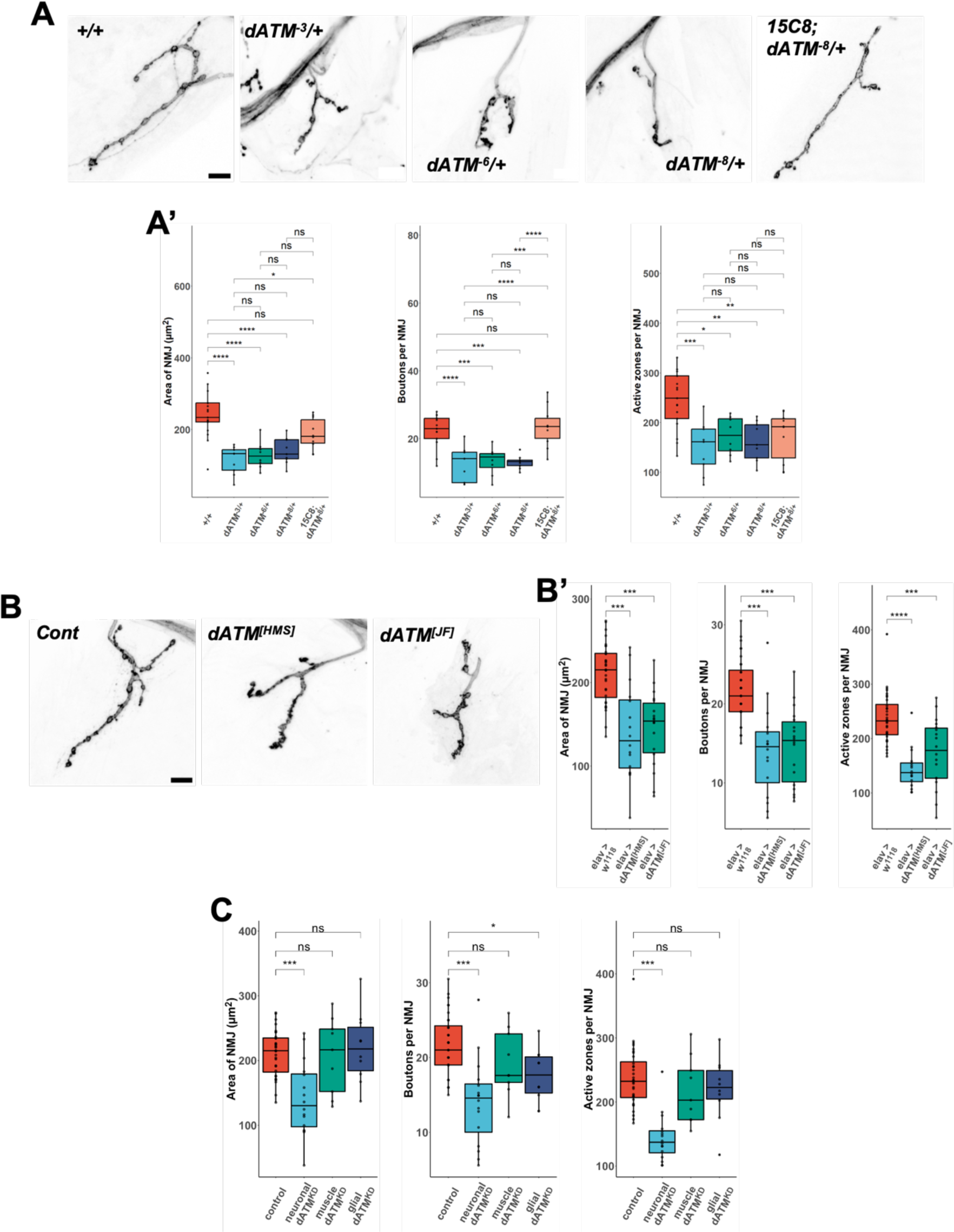
Presynaptic *Drosophila* ATM (dATM) is required for NMJ growth. (A) Representative images of NMJ4b of the indicated *dATM* genotypes stained with an--HRP to visualised the neuronal membrane. Graphs below the images show quantification of 3 metrics (NMJ surface area, bouton count and active zone count) of control *vs.* 3 different heterozygous *dATM* null alleles plus a genomic rescue construct: *15C8;dATM^-8/+^*, shown in pink. Tukey HSD test. (B) Representative images of NMJ4b showing control or Elav-Gal4 driving two different shRNA constructs for pre-synaptic knockdown of *dATM*. Quantification of NMJ metrics for each genotype shown below. Dunnett’s multiple comparisons test with *elav > w^1118^* as the reference group. (C) Quantification of NMJ features from a screen of presynaptic (neuronal), postsynaptic (muscle) and perisynaptic (glia) *dATM* knockdown. Dunnett’s multiple comparisons test with control as the reference group (*Gal4>w^1118^* for each driver). Individual data points are shown with boxes representing the median and interquartile range. p≤0.05 *, p≤0.01 **, p≤0.001 ***, p≤0.0001 ****, ns = not significant. Scale bars = 10 μm.

While confirming a requirement for neurodevelopment for dATM, these experiments do not address in which cell type dATM function is crucial for neurodevelopment, since the entire animal is heterozygous. The larval NMJ is composed of the presynaptic neuron, the postsynaptic muscle, and perisynaptic glia. One key advantage of *Drosophila* as a model for neurodevelopment is the ability to interrogate spatiotemporal requirements of genes-of-interest through the GAL4-UAS system. We used neuronal, glial or muscle-specific Gal4 drivers to express UAS-shRNAi to knockdown *dATM* in a cell-type specific manner. Neuronal knockdown using *elav-GAL4* with either of two different shRNA constructs phenocopied the *dATM* heterozygous phenotype, with marked reductions in NMJ surface area, bouton number, and active zone count (Fig 1B). In contrast, neither glial nor muscle knockdown of *dATM* had any effect on the morphology of the NMJ (Fig 1D). This strongly suggests the requirement for presynaptic dATM during *Drosophila* larval NMJ development.

To assess whether the structural developmental deficits we see have functional consequences for the neuron and for the animal, we recorded from the intersegmental neuron which innervates muscle 4 and, since the NMJs innervate the larval body wall muscles, we measured speed of crawling. Pan-neuronal *dATM* knockdown resulted in a significant decline in excitatory junction potential (EJP) and miniature EJP (mEJP) amplitudes (Fig 2A and 2B), while no overall change in mEJP frequency was observed (Fig 2C). There was also a small but significant decrease in quantal content (Fig 2D), and an increase in paired-pulse ratio (PPR, Fig 2F), likely due to a retention of the readily-releasable pool of synaptic vesicles caused by an overall reduction of the amplitudes of each EJP in the PPR test. Consistent with an electrophysiological deficit, *dATM* knockdown larvae crawled at reduced speed compared to controls (Fig 2H). There was, however, no overall change in behaviour of the knockdown larvae – they spend the same proportion of time moving as the controls (Fig 2I).

**Figure 2.**
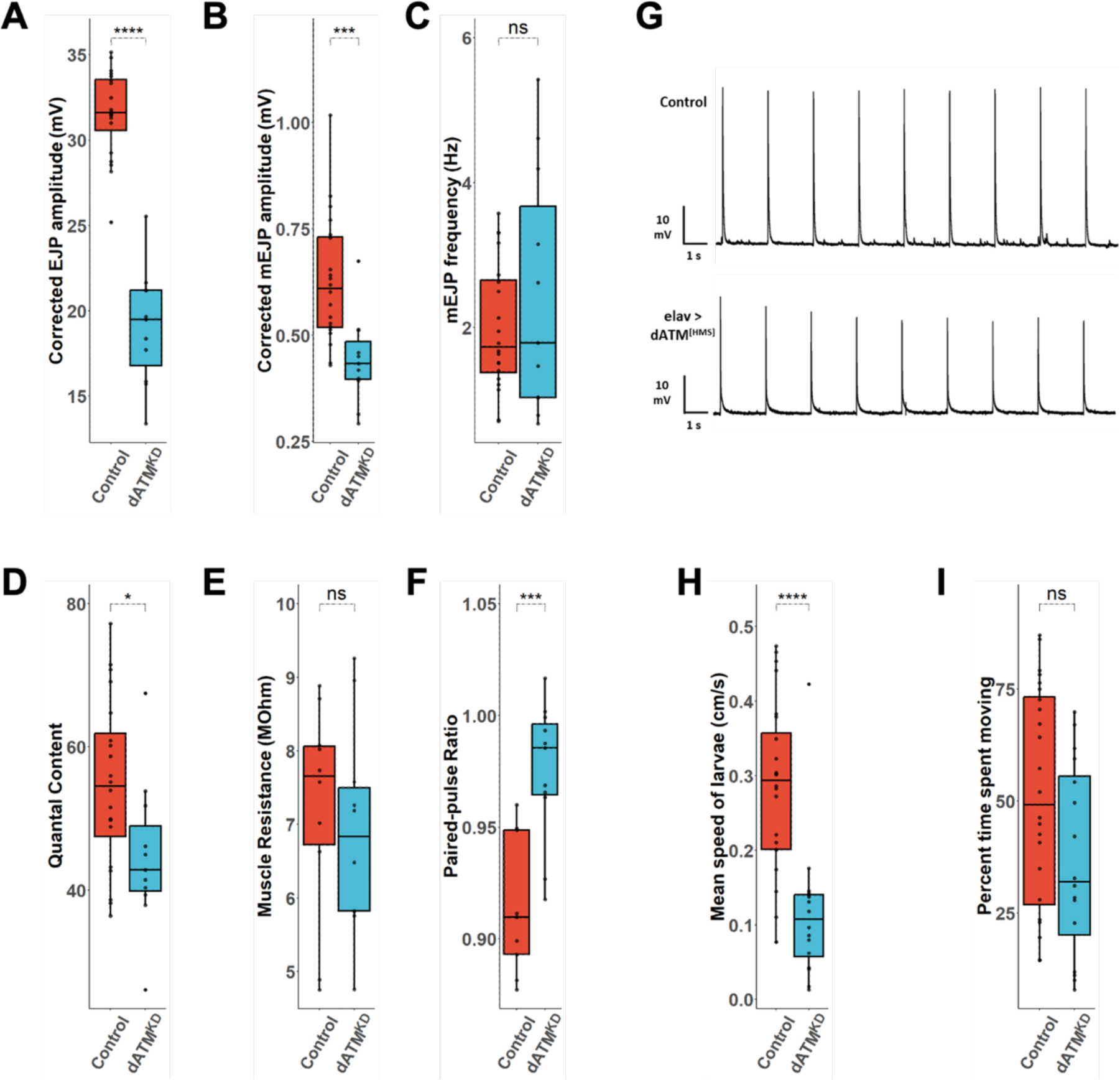
Presynaptic dATM is required for NMJ function and larval locomotion. (A-F) Quantification of electrophysiology parameters: (A) Corrected EJP amplitude; (B) Corrected mEJP amplitude; (C) mEJP frequency; (D) Quantal content; (E) Muscle resistance; (F) Paired-pulse ratio. (G) Representative electrophysiological traces of evoked EJPs from the indicated genotypes. (H-I) Quantification of larval locomotion parameters: (H) Mean larval crawling speed; (I) Mean percentage -me moving during experiment. In all plots, dATM^KD^ = presynaptic *dATM* knockdown. All p values from Student’s *T* tests with Welch’s correc-on, p≤0.05 *, p≤0.01 **, p≤0.001 ***, p≤0.0001 ****, ns = not significant.

### Neurons deficient in dATM fail to expand during development and show signs of degeneration at higher rearing temperatures

Neuronal activity plays a crucial role in shaping the formation and plasticity of connections in developing nervous systems, including the larval neuromuscular system. As *Drosophila* are poikilothermic, external temperature substantially affects their development, aging, and activity. Higher temperatures shorten their developmental period and enhance mobility, evident in both larval and adult stages. Consequently, there is an increase in neuronal activity with increased temperature, and a concomitant increase in NMJ size and arborisation (41).

To test the dependence of the neuronal *dATM* knockdown phenotype on developmental rearing temperature, presynaptic *dATM* knockdown was repeated with rearing temperatures of 19°C or 27°C (Fig 3A). Consistent with the work of others, e.g. (41,42) we observed that in the controls, increasing rearing temperature significantly increased the size and bouton number of NMJs, although active zone count appears to be unaffected by rearing temperature in our experiments. At both low and high rearing temperatures, presynaptic *dATM* knockdown results in significant deficits in NMJ development when compared to the controls: the knockdown larvae completely failed to expand their NMJs in response to increasing rearing temperature (Fig 3A). It appears that neuronal dATM may be required to sense the increased activity, act on that signal and respond appropriately by driving expansion.

**Figure 3.**
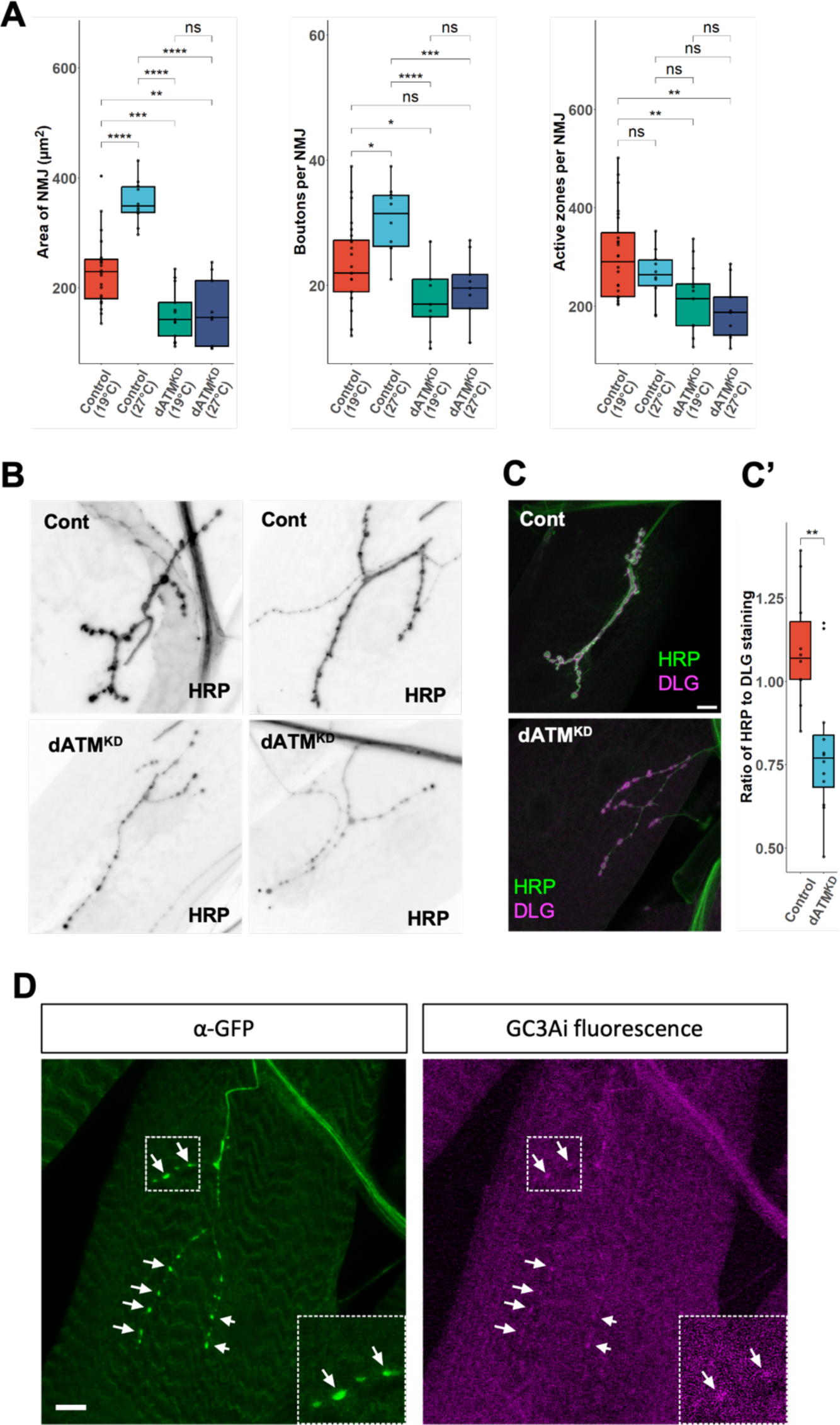
dATM-depleted neurons show signs of neurodegeneration at higher rearing temperatures. (A) Quantification of NMJ features in control *vs.* presynaptic *dATM* knockdown larvae at low (19°C) and high (27°C) rearing temperatures. Tukey HSD test. (B) Representative images of control *vs.* presynaptic *dATM* knockdown NMJs at 27°C rearing temperature. Note the thinning fragmentation of the presynaptic membrane as visualised by HRP staining. (C) Representative images of the same genotypes from B co-stained with presynaptic (HRP) and postsynaptic (DLG) markers. (C’) Quantification of the ratio of HRP to DLG staining from the indicated genotypes. P values from Student’s *T* tests with Welch’s correction. Individual data points are shown with boxes representing the median and interquartile range. p≤0.05 *, p≤0.01 **, p≤0.001 ***, p≤0.0001 ****, ns = not significant. (D) UAS*GC3Ai* expression in posterior motor neurons in presynaptic *dATM* knockdown larvae. *Le8 panel:* immunoreactivity from ⍺-GFP staining indicates expression of the GC3Ai reporter; *righ panel*: endogenous *GC3Ai* fluorescence indica-ng activated caspase. Boxed sec-on is shown in 2x zoom in the inset. Arrows indicate sites of localised caspase activity. Scale bars = 10 μm.

Higher activity levels in neurons places them under increased stress, particularly oxidative stress. Ataxia-telangiectasia features early-onset neurodegeneration, principally in the cerebellum which causes the ataxia in children. We were interested to see if depleting *dATM* in neurons would destabilise them at higher temperatures and lead to signs of degeneration.

A marker of neurodegeneration in the third instar larval model is withdrawal of the presynaptic membrane from the postsynaptic density, which remains in the muscle as a synaptic “footprint” (43). We shifted the flies from to the usual rearing temperature of 25 °C to 27 °C and knocked down *dATM* using the OK371 driver which drives in glutamatergic neurons, including the motor neurons innervating the body wall muscles. In *dATM* knockdown animals reared at 27°C, there were no notable examples of DLG staining which completely lacked the cognate HRP signal. However, there was a clear reduction in intensity of the pre-synaptic HRP signal corresponding to the neuronal membrane (Fig. 3B,C) and a clear decrease in the ratio of pre to postsynaptic signal in *dATM* knockdown neurons (Fig. 3C’), potentially an early indication of neurodegeneration.

We looked for localised caspase activity in the motor neuron terminals, which might underpin retraction of the neuron. We expressed the GC3Ai system in motor neurons as a reporter of (44). GC3Ai utilizes GFP molecules connected at the C- and N-termini by a DEVD caspase cleavage site. Without caspase activity, the linker keeps GFP fluorescence suppressed, though the protein can still be localised with GFP antibodies. When activated, caspases cleave the linker and GFP fluorescence is restored. Endogenous GFP activity is therefore indicative of caspase activity. In the long motor neurons innervating the posterior segments, endogenous GFP fluorescence was seen in boutons of the NMJ when *dATM* was knocked down, but not in controls (Fig. 3D; controls shown in supplementary results). Altogether, these data suggest that, at higher rearing temperatures, *dATM* knockdown neurons are on the cusp of neurodegeneration.

### Localisation of neuronal GFP-dATM

In mammals, neuronal ATM often shows strong cytosolic localisation and has been detected in synapses, colocalising with markers of presynaptic vesicles (20,45). Previous studies looking at dATM localisation overexpressed a FLAG-dATM construct in *Drosophila* S2 cells, revealing a predominantly nuclear localisation and foci formation upon irradiation (46). Given this study was restricted to *in vitro* overexpression in a non-neuronal cell type, the specific neuronal localisation of dATM in *Drosophila* remains unclear.

To address this gap in knowledge, a msGFP2-tagged full-length *dATM* cDNA was synthesised and cloned into pUAST-attB. msGFP2 was selected for its stability in oxidising conditions and the presence of point mutations in its dimerization interface (47). This latter point was crucial since if dATM’s function is orthologous to hATM, it may have differing downstream targets based on its dimerization state. Thus, it was essential to prevent artificial dimerization caused by GFP-GFP interaction.

*UAS-dATM[msGFP2]* expression was driven in glutamatergic neurons with *OK371-GAL4*. GFP fluorescence was visible in both the CNS and salivary glands (where *OK371-GAL4* drives off-target expression). Detailed examination revealed strong GFP expression in midline motor neuron cell bodies, peripheral motor neurons, and extending along axons from the ventral nerve cord (Fig. 4A). High magnification confirmed that dATM[msGFP2] expression is predominantly cytosolic in the motor neuron cell bodies (Fig. 4B). However, irradiation of larvae with X-irradiation which generates double-strand breaks in the DNA leads to a relocalization of dATM[msGFP2] into the nucleus, consistent with the role of ATM in the DDR (Fig. 4B).

**Figure 4.**
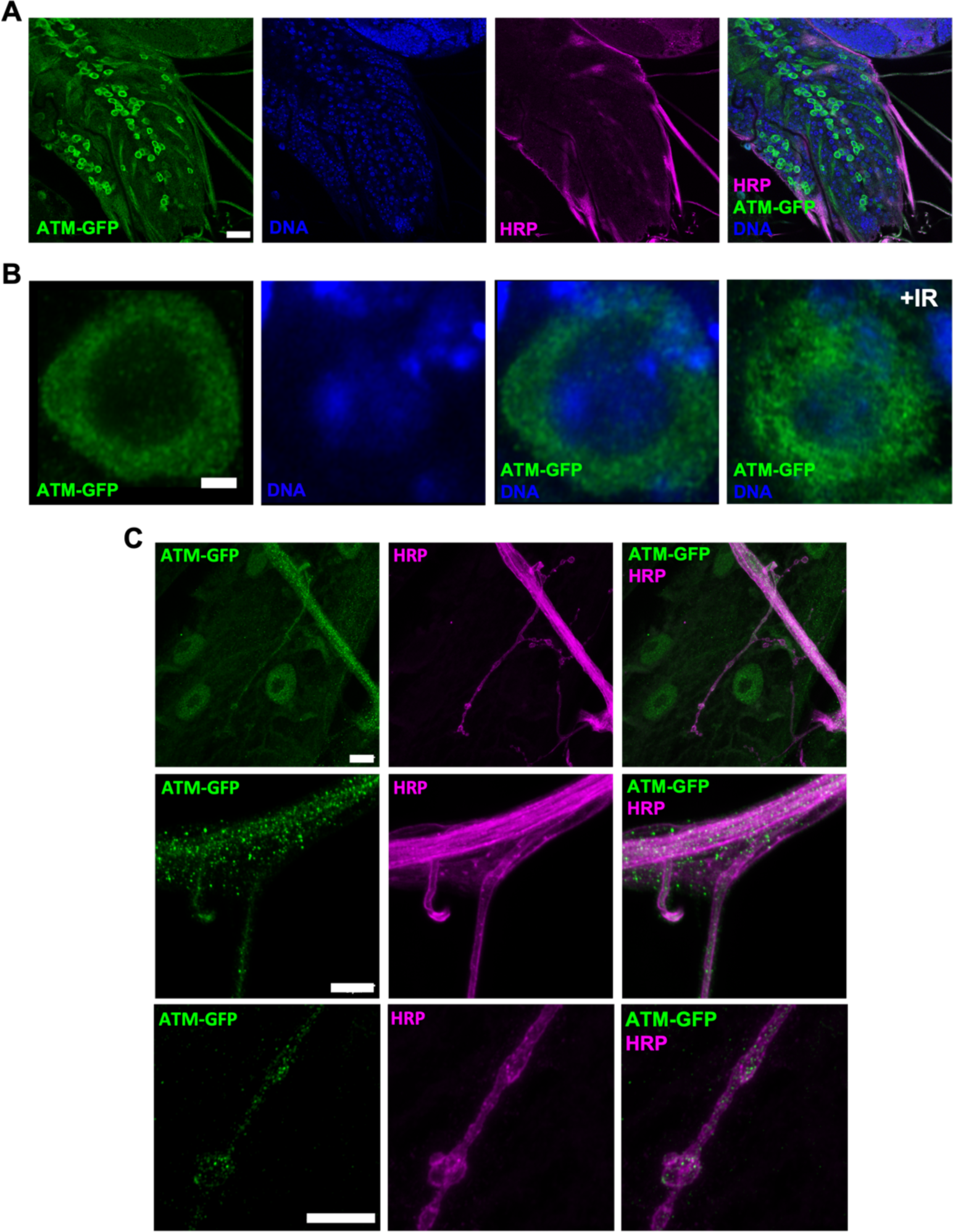
Motor neuron expression of msGFP2-tagged dATM. (A) Larval ventral nerve cord showing UAS-dATM[msGFP2] expression driven in glutamatergic neurons by *OK371-GAL4*. Scale bar = 20 μm. (B) Higher magnification of a single motor neuron cell body where dATM[msGFP2] expression is predominantly cytosolic under basal conditions but some relocates to the nucleus aker irradiation with 8 Gy of X-ray irradiation (right). Scale bar = 1 μm. (C) dATM[msGFP2] expression becomes increasingly punctate at regions distal to the motor neuron cell body: *Top –* lower magnification image of entire muscle 4 NMJ of *OK371-GAL4* driven dATM[msGFP2] expression, scale bar = 10 μm; *Middle –* higher magnification of axon, scale bar = 5 μm; *Lower –* higher magnification image of NMJ terminal, puncta of GFP appear to be concentrated within boutons (arrowheads). Scale bar = 5 μm.

In the periphery, OK371-driven dATM[msGFP2] expression appears more punctate than diffuse (Fig. 4C). The axon exhibits bright dATM[msGFP2] puncta, which become more pronounced in distal segments compared to regions proximal to the VNC. There also appears to be foci of dATM[msGFP2] external to the HRP stain of the axon, potentially indicating shuttling into the ensheathing glia. At the NMJ, presynaptic boutons display a notably higher density of msGFP2 puncta than the inter-bouton space (Fig. 4C, white arrowheads). Interestingly, low-level, punctate fluorescence was seen in the large nuclei of the muscle cells, possibly from some leaky expression. Taken together, neuronal dATM appears primarily cytosolic and localises to axonal and presynaptic puncta.

### The neurodevelopmental role of presynaptic dATM is independent of the DNA damage response

Given ATM’s key role in the DNA damage response (DDR) to double-stranded DNA breaks (DSBs), we hypothesized that the NMJ phenotype from presynaptic dATM knockdowns might stem from general DDR misregulation, rather than a specific involvement of dATM in neurodeveloment *per se*. To explore this, the expression of two key components of the DDR upstream and downstream of ATM was knocked down in neurons using the pan neural *elav-GAL4* driver to express a TRiP dsRNA construct. Specifically, the *Drosophila* homologs of the MRN complex component, MRE11, and the downstream checkpoint kinase 2 (CHK2, *loki* in *Drosophila*) were targeted. However, no detectable changes in the NMJ’s surface area or active zone count were observed compared to controls (Fig 5A. and 5B).

**Figure 5.**
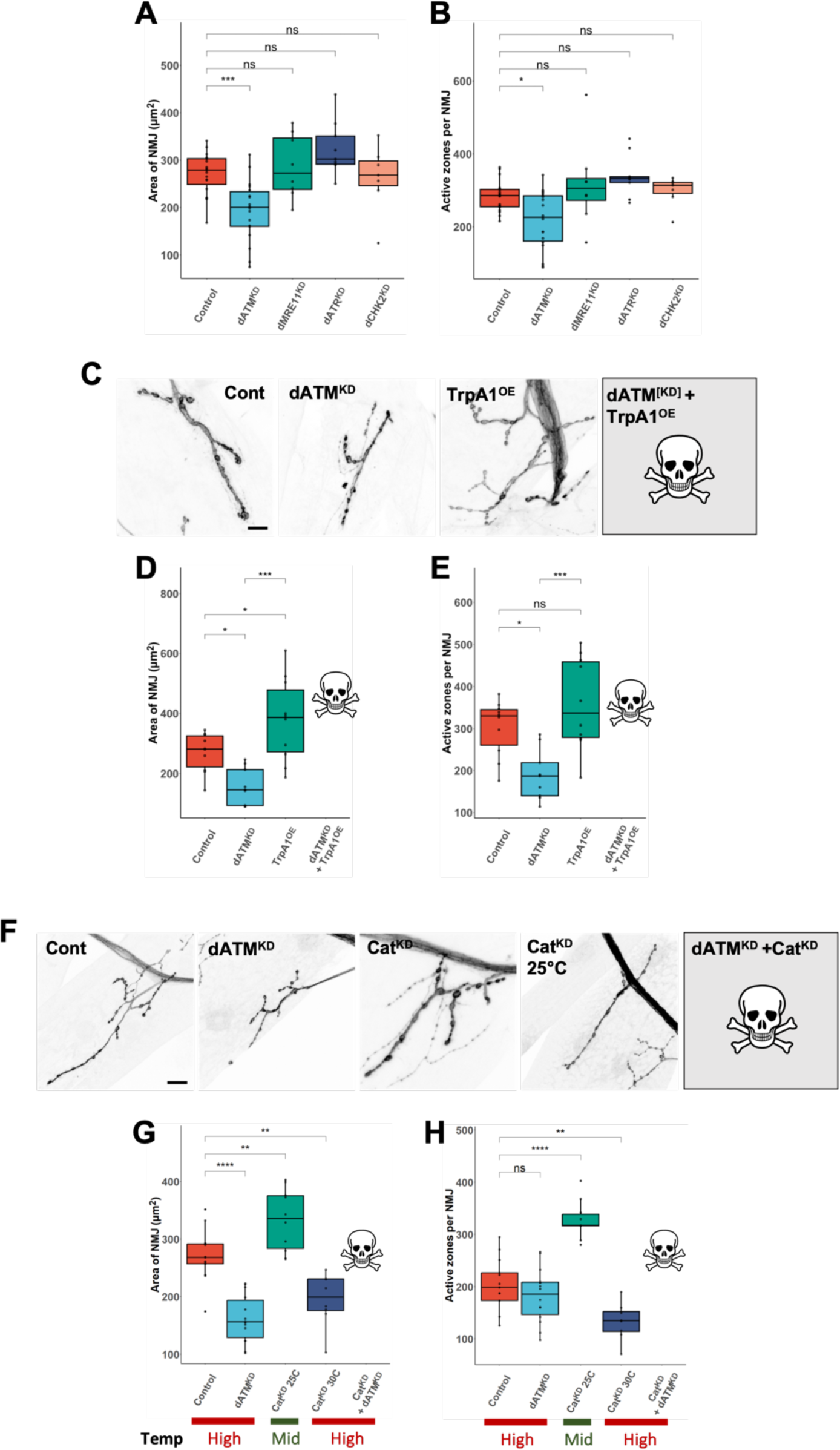
(A-B) Presynaptic knockdown of other DNA damage response components has no effect on the structural development of the NMJ as measured by (A) NMJ surface area or (B) active zone count. Dunnett’s multiple comparisons test with *Control* as the reference group. (C-E) Presynaptic overexpression of the temperature-gated ca-on channel *TrpA1* leads to expansion of the NMJ but is lethal in combination with *dATM* knockdown at 27°C rearing temperature. (C) representative muscle 4 NMJ images of the indicated genotypes; (D) NMJ surface area Quantification; (E) active zone count. Tukey HSD test. (F-H) Decreased ROS scavenging through *catalase* knockdown leads to NMJ expansion at moderate rearing temperatures (25°C) and reduced NMJ growth at high rearing temperatures (30°C), where it is lethal in combination with *dATM* knockdown. (F) representative muscle 4 NMJ images of the indicated genotypes; (G) NMJ surface area; (H) active zone count. Dunnett’s multiple comparisons test with *Control* as the reference group. p≤0.05 *, p≤0.01 **, p≤0.001 ***, p≤0.0001 ****, ns = not significant. All scale bars = 10 μm.

Further, since there is evidence elsewhere showing an interaction between neuronal ATM and its sister kinase, ATR, at synapses, we wanted to ask whether knockdown of *dATR* (*mei-41* in Drosophila) would replicate the effect of dATM knockdown. However, as with the other DDR components, we observed no significant difference in NMJ surface area or active zone count compared to controls (Fig. 5A and 5B). Given that targeting of upstream, downstream, or parallel components of dATM do nor replicate the effect of *dATM* knockdown, and that the dATM protein is primarily cytosolic in neurons, it seems likely that the role for dATM in neurodevelopment is independent of its role in the DDR. This suggested to us that key to understanding the developmental role of presynaptic dATM lay with its cytosolic pathways and interactions.

### Presynaptic dATM knockdown sensitises larvae to excitotoxicity and oxidative stress

In addition to its role in the DDR, ATM is a key player in oxidative stress signalling. Specifically, it is known that cytosolic, dimeric ATM can be directly oxidised by ROS, leading to an intermolecular disulphide bond forming at Cys 2991 and conversion of the dimer into an active state, with distinct downstream targets outside of the DDR (14,48). Given the signs of early degeneration in *dATM-*depleted neurons, we considered that this may be caused by impaired oxidative stress signalling.

In *Drosophila*, there is increasing evidence that ROS signalling regulates larval NMJ development and plasticity. For example, *spinster* mutants display elevated ROS levels and expanded NMJs, which can be rescued through increased ROS scavenging (49). Expression of the temperature-gated cation channel, TrpA1, with a rearing temperature ≥25°C hyper-activates neurons and results in increased mitochondrial ROS and consequent NMJ overgrowth. Consistent with this, reducing antioxidant capability of neurons through catalase knockdown phenocopies this effect (1). However, excessive oxidative stress can lead to neurodegeneration (50) and thus a delicate balance must be maintained.

We hypothesised that the *dATM* knockdown phenotype could be a failure to sense normal changes in ROS or activity levels during NMJ maturation, resulting in a failure for the NMJ to expand appropriately. This could potentially be overcome through hyperactivating the neuron with TrpA1 throughout development to compensate. We combined knockdown of *dATM* in motor neurons with overexpression of the TrpA1 and reared the larvae at 27°C, which results in tonic neuronal firing (51). In larvae overexpressing TrpA1 without *dATM* knockdown, this produced a significant expansion of the NMJ (Fig. 5D), as reported previously (1) although no changes to active zone count were observed (Fig. 5E). In marked contrast, co-expression of TrpA1 with *dATM* knockdown was lethal before late larval stage, suggesting that *dATM* knockdown sensitises neurons to excitotoxicity.

To explore whether this was the result of an increased ROS burden, we combined *dATM* and *catalase* knockdown. As reported previously, *catalase* knockdown at the standard 25 °C rearing temperature leads to an expansion of the NMJ due to the decreased ROS scavenging capacity (Fig. 5G,H). However, raising the temperature to 30 °C to increase neuronal activity results in a significant undergrowth of the NMJ (Fig. 5G,H), potentially because the combination of increased ROS from activity and reduced scavenging passes the threshold for toxicity. Here, as with *TrpA1* overexpression, the combination of *catalase* and *dATM* knockdown is lethal, indicating that *dATM-* depleted neurons cannot cope with a reduction in the ROS-scavenging machinery. Taken together, these results indicate that *dATM-*depleted neurons are hypersensitive to oxidative stress.

### Presynaptic dATM interacts with the autophagy machinery in NMJ development

We noted that the *dATM* null heterozygotes and neuronal knockdowns phenocopied mutations and knockdowns of the autophagy machinery, which is itself another key regulator of NMJ development in *Drosophila; atg1* mutants show significant underdevelopment of the NMJ, while synaptic overgrowth can be induced by overexpressing ATG1 in neurons (4). We had also noted that the localization of dATM[msGFP2] in neurons i.e. a diffuse cytosolic localization in the soma with discrete punctate along the axon and extending into the synapse, is consistent with the known localisation of neuronal autophagosomes. These typically form distal to the soma in axons and are transported retrogradely along the axons by dynein motors (52,53). Given the established role of cytosolic ATM linking oxidative stress signalling to autophagy (15,16), we therefore wanted to investigate whether dATM knockdowns would have alterations in autophagic flux, and whether any interaction with the autophagy machinery would be observed.

To quantify macroautophagic flux, we utilised the tandem *GFP-mCherry::Atg8a* “traffic light” reporter, overexpressed in glutamatergic neurons using the *OK371-GAL4* driver. This reporter relies on the relative pH sensitivities of its constituent fluorescent proteins. *Atg8a* localises to autophagosomes, where both GFP and mCherry are fluorescent, forming yellow puncta. Upon autophagosome maturation into autolysosomes, the acidic environment results in quenching of the GFP signal, while mCherry fluorescence is maintained, resulting in red puncta (34,54).

To induce autophagy, feeding-stage third instar larvae were removed from the food and starved of amino acids for 4 h in a 20% sucrose solution before the ventral nerve cords were dissected for live imaging. Surprisingly the GFP signal remained diffuse although the mCherry signal was punctate, as expected (Fig. 6A). We used a custom FIJI routine to measure the intensity of mCherry fluorescence in puncta plus the ratio of GFP:mCherry fluorescence in each. This reported a lower intensity of mCherry fluorescence in the *dATM* knockdown neurons (Fig. 6B) and a considerably higher ratio of GFP:mCherry, which we interpret to represent a failure of the autophagosomes to mature into autolysosomes (Fig. 6B).

**Figure 6.**
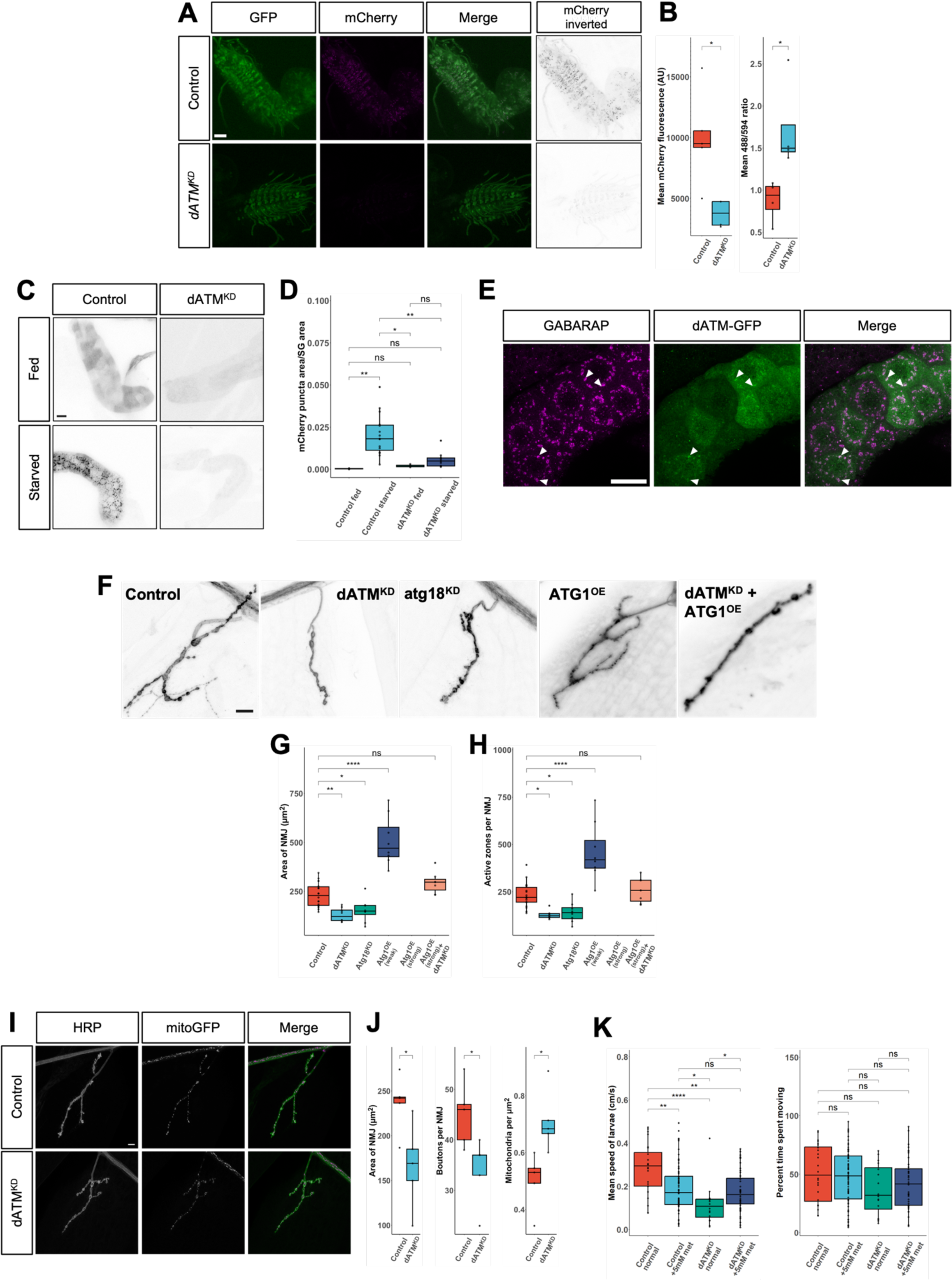
The interaction of dATM with autophagy. (A,B) Expression of the tandem *GFP-mCherry::Atg8* reporter of autophagic flux in feeding-stage larvae starved for 4 h. The reporter is expressed in glutamatergic neurons of the ventral nerve cord under the control of *OK371-Gal4* and imaged live. GFP fluorescence reports early autophagosomes but remains diffuse. mCherry reports both autophagosomes and low pH late autophagolysosomes and is punctate. The fluorescence intensity of mCherry puncta is significantly reduced by knockdown of *dATM* (B) and the ratio of GFP (488 mCherry (594) fluorescence is significantly increased (B). (C,D) Localisation of the mCherry fluorescence of the tandem reporter expressed and imaged live in salivary gland cells. mCherry fluorescence is diffuse in salivary gland cells from feeding-stage larvae (C, upper panels). After 4 h starvation mCherry+ puncta are present in Control cells representing induction of autophagy but not in *dATM* knockdown cells where fluorescence remains diffuse (C, lower panels). mCherry+ puncta are quantified relative to area of the salivary gland cells in (D). (E) dATM-sfGFP (green) co-localises with autophagosomes labelled with anti-GABARAP (magenta) in starved salivary gland cells. Arrowheads point to examples of colocalized foci. (F-H) Genetic interactions between *dATM* and components of the autophagy machinery. Knockdown of *atg18* leads to undergrowth of the NMJ and phenocopies knockdown of *dATM.* Overexpression of *ATG1* leads to overgrowth but concurrent overexpression of *ATG1* with knockdown of *dATM* restores the NMJ to the size of Controls. (F) Representative muscle 4 NMJ images of the indicated genotypes; (G) NMJ surface area; (H) active zone count. (I,J) Mitochondrial density increases in the NMJ after *dATM* knockdown. (I) Representative images of NMJ4 of Control (upper row) and *dATM* knockdown (lower row) expressing the mitoGFP reporter then stained for HRP to visualize the neuronal membrane and GFP for mitochondria. (J) Quantification of NMJ surface area, boutons number per NMJ and the density of mitochondria per μm^2^ of NMJ surface. (K) Supplementation of food with 5 mM metformin which induces autophagy via activation of AMP kinase significantly reduces the locomotive speed of Control larvae but increases the speed of *dATM* knockdown larvae to the same speed as Controls. Percentage time larvae spend moving is not affected by metformin supplementation. p<0.05 *, p≤0.01 **, p≤0.001 ***, p≤0.0001 ****, ns = not significant. Scale bars = 40 μm (A), 30 μm (C), 40 μm (E) and 10 μm (I).

Imaging of autophagosomes in the CNS neurons proved difficult so for confirmation of an autophagy deficit, we took advantage of the off-target driving of the Atg8 traffic light reporter by *OK371-GAL4* in the salivary glands. In the fed state, both control and *dATM* knockdown salivary glands show diffuse GFP and mCherry signals. After 4 h starvation, significant numbers of mCherry+ puncta were visible in the control salivary gland cells, indicating increased autophagic flux. However, these were conspicuously absent from *dATM* knockdown cells (Fig. 6C,D) confirming that *dATM* knockdown cells are unable to induce autophagic flux in response to starvation.

We next asked whether dATM may be present in autophagosomes. We expressed *dATM[msGFP2]* with *OK371-GAL4* and starved the larvae for 4 h, as before. We could see detect clear colocalization of GFP-dATM with the autophagosome marker, anti-GABARAP (Fig. 6E) in the salivary gland cells.

Our results pointed to a requirement for dATM for the induction of autophagy but we sought to strengthen this conclusion by looking for genetic interactions between *dATM* and the autophagy machinery. Consistent with a previous study of autophagy in NMJ development (4), we found that presynaptic knockdown of *Atg18* phenocopies presynaptic *dATM* knockdown, and that overexpression of ATG1 from a weaker UAS-line leads to significant synapse expansion (Fig. 6F,G). However, strong overexpression of ATG1 is lethal: as with oxidative stress, the levels of autophagy appears to be held in a delicate balance during synapse development. Significantly, combining *dATM* knockdown with strong ATG1 overexpression is no longer lethal and rescues the synapse development deficits of *dATM* knockdown larvae (Fig. 6F,H). These data are a clear indication of an interaction between *dATM* and the induction of the autophagy machinery.

ATM has been reported previously to stimulate mitophagy via PINK1/Parkin (55–57). We asked whether mitophagy was affected in motor neurons by knockdown of *dATM* driving expression of the reporter, UAS-mitoGFP, concurrently with shRNA to *dATM*. Mitochondria in the NMJ were then counted using a FIJI routine. Knockdown of *dATM* resulted in a significant increase in mitochondrial density, indicating that mitophagy in these neurons may indeed be defective (Fig. 6I,J). A failure in mitophagy has the potential to increase oxidative stress since defective mitochondria are not recycled. This may contribute to the susceptibility of the *dATM* knockdown neurons to increased ROS, seen earlier (Fig. 5).

Finally, we asked whether chemical induction of autophagy could rescue the locomotor deficit exhibited by the *dATM* knockdown larvae. We supplemented food with 5 mM metformin, a potent inducer of macroautophagy thought to function via activation of AMP kinase (58). Interestingly, metformin supplementation significantly diminished locomotor function of control larvae compared to larvae raised on standard food (Fig. 6K), However, metformin was beneficial to the locomotor performance of presynaptic *dATM* knockdown larvae whose crawling speed was raised to the same level as the metformin-fed control larvae. (Fig. 6K). These results further support the notion that autophagy levels must be held in delicate balance and that presynaptic *dATM* is required for neurons to maintain this balance.

### AMP kinase acts downstream of pre-synaptic dATM to induce autophagy

If dATM is acting downstream of an oxidative stress signal to regulate autophagic flux and expand the synapse, we reasoned that this may be occurring through the canonical ROS-ATM-AMP kinase (AMPK) signalling axis previously identified in mammalian cell studies (15). We overexpressed or knocked down *AMPK* in neurons with, or without, concurrent *dATM* knockdown. Interestingly, *AMPK* knockdown alone did not significantly alter NMJ development (Fig. 7A-C) and the combination of *AMPK* and *dATM* knockdown phenocopied the of *dATM* knockdown alone and the NMJs failed to expand. There was, however, a marked fragmentation of the presynaptic membrane in double knockdown NMJs with occasional bright spots, reminiscent of “retraction bulbs” seen with degenerating neurons.

**Figure 7.**
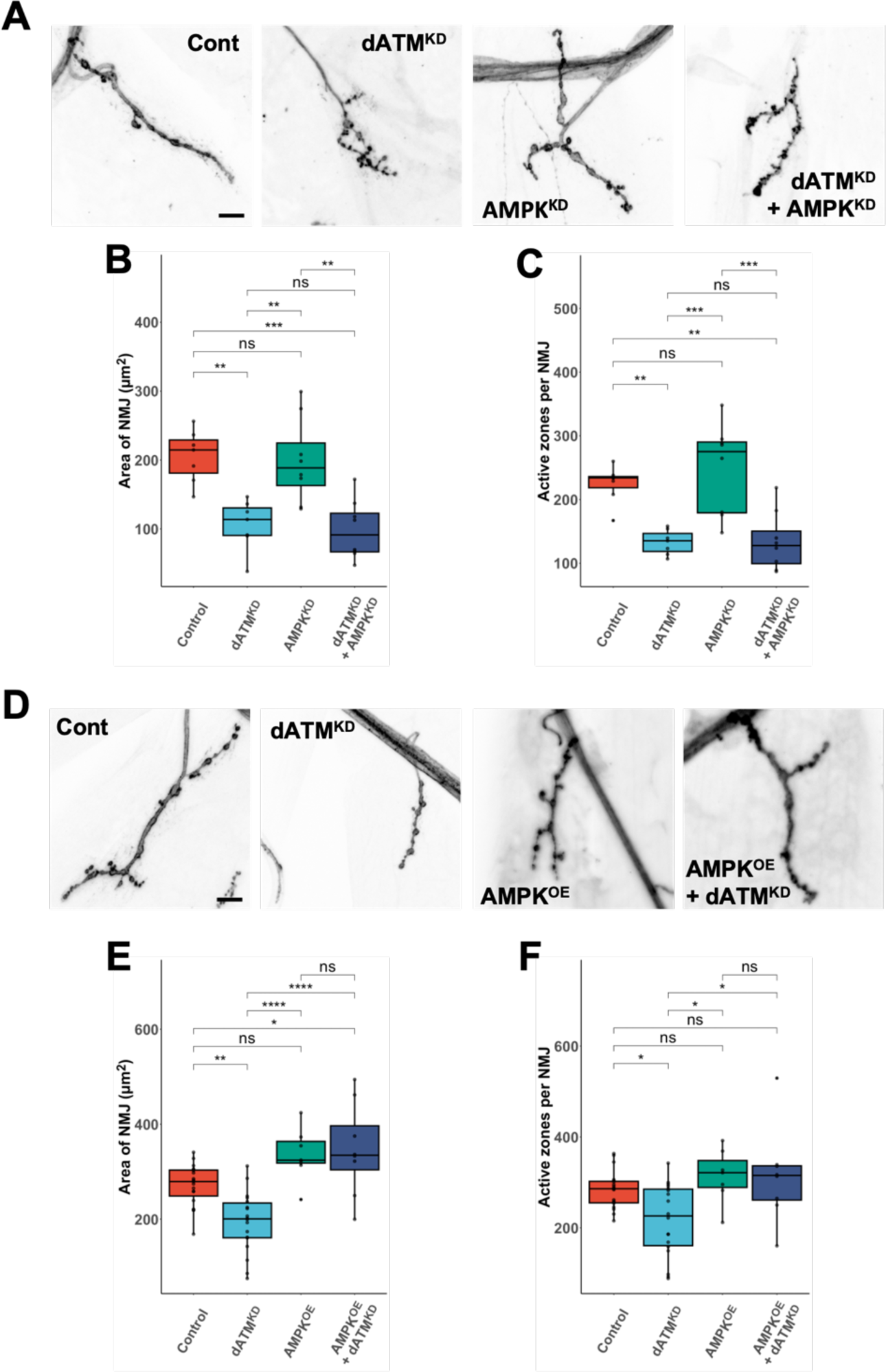
AMP kinase acts downstream of dATM to regulate NMJ development. (A-C) Single and concurrent presynaptic knockdowns of *dATM* and *AMPK*. Presynaptic *AMPK* knockdown alone has no impact on NMJ structural development. Concurrent knockdown of *AMPK* with *datum* phenocopies single *dATM* knockdown: (A) Representative muscle 4 NMJ images of the indicated genotypes; (B) NMJ surface area; (C) active zone count. Tukey HSD test. (D-F) Overexpression of *AMPK* singly and concurrently with *dATM* knockdown. Overexpression of *AMPK* has a non-significant effect on NMJ development. In combination with *dATM* knockdown NMJs expand significantly: (D) Representative muscle 4 NMJ images of the indicated genotypes; (E) NMJ surface area; (F) active zone count. Tukey HSD test. p≤0.05 *, p≤0.01 **, p≤0.001 ***, p≤0.0001 ****, ns = not significant. Scale bars = 10 μm.

AMPK overexpression resulted in a small but non-significant increase in NMJ size (Fig. 7D-F) and overexpression combined with *dATM* knockdown rescue the undergrowth phenoptype and expanded the synapse to a significant degree vs. controls (Fig. 7D-F). These genetic epistasis experiments are consistent with dATM acting through AMPK to induce autophagy in response to increasing ROS levels in NMJ development, thereby regulating synapse expansion.

## Discussion

Mutations in ATM kinase result in the progressive early-onset neurodegenerative disorder ataxia-telangiectasia (A-T), although the underlying disease mechanism is not well understood (59,60). ATM has well-characterized nuclear functions in the DNA damage response but there is debate around the extent to which neuronal ATM is primarily cytosolic or nuclear, and thus which of its nuclear *vs.* cytosolic functions are most relevant to the vulnerability of neurons in A-T (18,19,61). In addition, there is an increasing understanding that ATM may have unique roles in neurons compared to other cell types (20,45,62). Here, we have demonstrated that the *Drosophila* homologue, *dATM*, is required specifically presynaptically for normal synapse development, function and homeostasis.

The structural and functional phenotypes of both *dATM* null heterozygosity and presynaptic *dATM* knockdown are indicative either of an under-grown synapse which has failed to respond to growth signals, or of a synapse in the early stages of degeneration. The failure of *dATM*-deficient motor neurons to expand in response to increases in developmental rearing temperature suggests a failure of the neuron to transduce activity-dependent growth signals (42). However, these larvae were also vulnerable to artificial chronic neuronal over-activation or a reduction in antioxidant protection. Further, at higher rearing temperatures, there were indications of presynaptic retraction from the postsynaptic density, and local caspase activity within boutons, suggesting that these neurons are also on the cusp of degeneration.

As a neuron matures, it continually processes signals from numerous interconnected pathways, influencing its synaptic connectivity, strength, and structure. Our study has concentrated on the processes of autophagy and oxidative stress signalling; both pathways are known to positively regulate synapse expansion in *Drosophila*, yet when overly active, are detrimental to the health of neurons (1,4,49). Similarly, neuronal activity is essential for normal synapse development and maintenance, but excessive activity leads to excitotoxicity (41,42) which highlights the delicate balance the neuron faces during development and homeostasis. Our findings suggest that neuronal *dATM* plays a key role in transducing ROS-autophagy signalling and in maintaining a balance between these interconnected processes.

For instance, presynaptic *dATM* depletion sensitized larvae to excitotoxicity and decreased ROS-scavenging yet was protective against chronic autophagy upregulation. When autophagy was pharmacologically induced in control larvae, their locomotor performance declined, but this intervention proved beneficial for larvae with presynaptic *dATM* knockdown. This aligns with other *Drosophila* research, both in neuronal and non-neuronal tissues, indicating an optimal level of autophagy in promoting lifespan and health (63,64). This finding also correlates with broader mammalian studies underscoring the importance of balanced autophagy for maintaining neuronal health (6). Enhanced autophagy has been shown to help clear toxic proteins that aggregate in disorders like Parkinson’s (65) and Alzheimer’s disease (66). However, excessive autophagy can itself result in neurodegeneration in different contexts (5,67).

In our model, presynaptic ATM responds to local ROS production generated through neuronal activity by activating the autophagic machinery through the conserved ATM-AMPK axis (15): an increase in neuronal activity stimulates expansion, a reduction in activity causes the reverse (4,41). This pathway may then be held in balance with other redox-sensitive ATM pathways, such as p53-mediated pro-apoptotic signalling, upregulation of mitophagy through Parkin, or promotion of the pentose phosphate pathway (PPP) to buffer oxidative stress. If ATM is a nexus for redox signalling, this would explain why *dATM*-depleted neurons are vulnerable to excitotoxicity or decreased ROS scavenging. Other work has shown redox-activated mammalian ATM upregulates the PPP via Hsp70 phosphorylation, increasing antioxidant capability (68,69), so this is a potential mechanism for how ATM activation at synapses could provide local homeostatic feedback to buffer ROS levels and prevent toxicity. Further epistasis work overexpressing or knocking down *Drosophila* PPP components may help to elucidate the precise nature of this feedback mechanism.

There is some controversy about the role of autophagy in the aetiology of the A-T. For example, neuronal precursor cells derived from A-T patients exhibit impaired autophagic flux and disrupted mitophagy (23), while pharmacological inhibition of autophagy rescued survival and synapse loss of ATM-deficient mouse cortical neurons (22). Given that the mouse A-T model lacks cerebellar degeneration, the fact that our *Drosophila* model recapitulates the deficient autophagic flux of human A-T neuronal precursors and shows evidence of neurological alterations suggest that it could prove to be a useful screening tool to identify potential treatments that stimulate autophagic flux.

While other studies have used *Drosophila* to examine the effects of *dATM* mutations and knockdowns on the structure of the adult brain, adult locomotion, and lifespan (70,71), this is the first to investigate the consequences on neurodevelopment and function. We believe this approach shows greater relevance to the progression of A-T, given its early-onset nature (12) and evidence of developmental patterning defects in the cerebellar architecture (72). With adult flies, there is a risk of failing to dissociate between the role of *dATM* in neural progenitors or in coordinating some aspect of neurodevelopment in metamorphosis, which would be distinct from mammalian ATM. Inconsistency in the literature exists as to the specific consequence of neuronal knockdown of *dATM*: some studies describe photoreceptor degeneration and temperature-dependent lethality (71); others report that it is glia which are vulnerable to *dATM*-deficiency and not neurons (40). Recent work in our lab has demonstrated that limiting *dATM* knockdown to adult neurons was neuroprotective and extended lifespan in different *Drosophila* models of neurodegenerative disorders (73). It seems likely that ATM plays different roles in cycling *vs.* non-cycling cells of the nervous system and in developing *vs.* matured neurons.

We found that neuronal knockdown of other DDR components did not recapitulate the *dATM* knockdown phenotype. While mutations in DDR proteins, such as components of the MRE11-Rad50-NBS1 complex, are associated with microcephaly (74–76) and neurodegeneration (77,78), a probable mechanism for the pathology in these DDR-related conditions is the death of neuronal precursors. This should not be a factor in our experiments; the shRNA to each component is almost exclusively being expressed in differentiated, post-mitotic neurons. Clearly there is a dissociation of the necessity of different DDR proteins in neuronal precursors *vs.* differentiated neurons, especially given our recent findings that knockdown of DDR components in a mature nervous system can be neuroprotective (38,73).

Neurons expressing *dATM[msGFP2]* showed a predominantly cytosolic localization of GFP. which became more nuclear only after DNA damage was induced. This localization, coupled with the epistatic interactions of ATM with the autophagy machinery and AMPK we demonstrate here, supports the idea that the extranuclear, redox-dependent signalling pathways of ATM are critical for its functions in neurons.

We have a growing understanding of the unique role for cytosolic ATM in neurons, separable from the DDR. This includes the physical association of ATM with synaptic vesicle proteins VAMP2 and synapsin-I (45), the regulation of excitatory *vs.* inhibitory neurotransmitter release (62), its role in LTP (20), and association with mitochondria and concomitant regulation of mitophagy (79). Our results show that cytosolic ATM is critical for neurodevelopment, acting to regulate the homeostatic expansion of synapses in response to changes in activity.

## Acknowledgements

The authors would like to thank Dr Ioannis Nezis for sharing the GFP-mCherry-Atg8 reporter line and for advice on visualising autophagy and the West Midlands *Drosophila* community for support and advice. MJT was funded by the Biotechnology and Biological Sciences Research Council Midlands Integrative Biosciences Training Partnership.

